# Sensitivity to Immune Checkpoint Blockade and Progression-Free Survival is associated with baseline CD8^+^ T cell clone size and cytotoxicity

**DOI:** 10.1101/2020.11.15.383786

**Authors:** RA Watson, O Tong, R Cooper, CA Taylor, PK Sharma, A Verge de Los Aires, EA Mahé, H Ruffieux, I Nassiri, MR Middleton, BP Fairfax

## Abstract

Immune checkpoint blockers (ICB) exert their anti-cancer effects via CD8^+^ T cells, although how responses vary over sub-populations and across clones is incompletely understood. We performed single-cell RNA-sequencing of CD8^+^ T cells and their receptors pre- and post-ICB across eight patients, integrating results with bulk-sequencing data (n=209). We identify seven subsets with divergent responses to ICB, finding the effector cluster demonstrates the most pronounced changes. Likewise, transcriptomic response to ICB relates to clone size, with large clones demonstrating increased numbers of regulated genes of higher immunological pertinence. Cytotoxic effector clones were more likely to persist long-term following ICB and overlapped with public tumour-infiltrating lymphocyte clonotypes. Notably, pre-treatment CD8^+^ cytotoxicity associated with progression-free survival, highlighting the importance of the baseline CD8^+^ immune landscape in long-term response. This work further advances understanding of the molecular determinants of ICB response and assists in the search for peripheral prognostic biomarkers.

**ONE SENTENCE SUMMARY:** Using single-cell and bulk RNA sequencing we explore checkpoint immunotherapy activity on peripheral CD8^+^ T cells in metastatic melanoma; demonstrating that cell subset and clone size determine gene expression responses to treatment, and that pre-treatment cytotoxicity and clonality of peripheral CD8^+^ T cells is clinically prognostic.

## INTRODUCTION

Antibodies binding the checkpoint proteins CTLA-4 (cytotoxic T-lymphocyte-associated protein-4) and PD-1 (programmed cell death protein-1) - Immune Checkpoint Blockers (ICB) - have revolutionised the treatment of metastatic melanoma (MM) and numerous other cancers*(1, 2)*. There is, however, marked heterogeneity in ICB-induced clinical benefit*(2)* and insights into determinants of ICB activity and variation in clinical response remain limited. Although tumour mutational burden has been linked to clinical outcome across multiple cancers*(3, 4)*, its absolute importance is unclear*(5)*. Similarly, the prognostically favourable association of tumour infiltrating lymphocytes (TILs) is long-established, but factors controlling T cell infiltration remain unresolved*(6, 7)*. Crucially, it is increasingly apparent that general aspects of the peripheral immune landscape may play a critical, yet comparatively under-investigated role in determining tumour recognition and control*(1, 2, 8)*.

CD8^+^ T cells are cardinal to the immune response towards MM*(8, 9)*, with most research focussing on their presence within TILs*(10, 11)*. However, recent analysis of non-melanoma skin cancer indicates the T cell response to ICB derives from a distinct repertoire of T cell clones, denoted by shared carriage of specific T cell receptors (TCR), that migrate to the tumour upon treatment*(12)*. In MM patients, T cell clones can be observed to be shared across tumour and blood compartments *(11, 13–17)*, whilst the number of expanded clones (occupying >0.5% of the repertoire) in the periphery post first cycle of ICB treatment associates with long-term clinical outcome*(18)*. Thus, there is robust evidence that the peripheral clonal T cell response to ICB is informative as to anti-cancer immune responses and may allow for on-treatment prognostication. Much of these analyses have been performed agnostic to CD8 T cell phenotypes, and the determinants of ICB activity at the level of subsets and on a clone-by-clone basis are unresolved. The CD8^+^ T cell ICB response has multiple components, including mitotic cell division*(19)* and increased cytotoxicity, as exemplified by interferon gamma (IFNγ) induction*(20)*, with the relationship between these being undefined. Interestingly, the magnitude of the mitotic response is not strongly linked to clinical outcome*(18)*. How these responses vary according to T cell phenotype*(21)*, their heterogeneity across clones, and if they are derived from novel or pre-existing clones is unknown. To explore this we have used single cell RNA-sequencing (scRNA-seq) to track the transcriptomic response of individual peripheral CD8^+^ T cell clones to ICB, identifying baseline markers of clonal sensitivity, and charting how response varies according to phenotypic subset and clonal size. We demonstrate that CD8^+^ T cell response to treatment with ICB varies according to phenotypic subset and clonal size, with the cytotoxic effector subset showing increased responsivity but remaining proportionally stable. Further dissection of ICB response at the clonal level reveals that clone size markedly impacts magnitude of response, with large effector T cell clones developing enhanced TCR signalling compared to other clones, and showing enrichment in melanoma TIL public clonotypes. By tracking TCRs over longer periods of treatment we demonstrate expansion and survival of pre-existing cytotoxic T cell clones, as well as markers of clonal persistence across cycles of treatment. Finally, we apply these observations regarding baseline clone size and cytotoxicity to a large clinical dataset, demonstrating that combining pre-treatment CD8^+^ cell cytotoxicity and large clone count is clinically prognostic, with individuals possessing both low cytotoxicity and fewer expanded clones typically gaining significantly less benefit from treatment with ICB.

## RESULTS

### Single-cell profiling of peripheral CD8^+^ T cells from patients with metastatic melanoma

We generated paired baseline (day 0 – ‘C1’) and on-treatment (day 21 – ‘C2’) 5’ scRNA-seq and V(D)J profiles across peripheral CD8^+^ T cells from eight patients with MM undergoing treatment with cICB (ipilimumab plus nivolumab, n=4) or sICB (pembrolizumab, n=4) (**fig. 1a; table S1; Methods**). In total, 22,445 *CD3/CD8* expressing cells passed quality control, of which 17,909 cells had TCR sequences. To identify distinct CD8^+^ T cell populations, we performed unsupervised clustering, observing globally similar UMAP projection of cells from both timepoints (**fig. S1a**), and subsequently cross-referenced each cluster to identities from published CD8^+^ T cell scRNA-seq datasets (**Methods**, **fig. S1b-e**), grouping cells aligned to each identity together. Seven distinct groups of CD8^+^ T cells were identified (**fig. 1b-c**); Naïve, central memory (CM), effector memory (EM) and effector cells (ECs) which collectively represent the majority of T cells, as well as highly cycling cells (Mitotic), γδ T cells (gd), and *KLRB1*-expressing mucosal-associated invariant T cells (MAIT), which formed a separate cluster from the rest of the cells. Mitotic cells had higher numbers of unique molecular identifiers (nUMI) and MKI67 expression per cell and a bias towards S and G2M cell cycle phases (**fig. S1f-g**), consistent with previous descriptions *(13, 22)*. Expression of *TRDC* was used to identify the γδ subset *(23)*, which was enriched for *KLRC3, KIR3DL2* and *KIR3DL3* (**fig. 1c**). To distinguish the biological features of each subset, we performed Gene Ontology Biological Process (GOBP) enrichment analysis (**fig. 1d**), which highlighted key immune pathways including *IFNG* signalling and T cell activation/co-stimulation were enriched in EM and ECs, consistent with higher cytotoxicity (**fig. S1h**). In contrast, Naïve/CM and Mitotic subsets demonstrated upregulated translational and mitochondrial machinery. We further examined the expression of T cell exhaustion markers across the cell subsets (**fig. 1e**). Increased expression of *TOX* and *HAVCR2* and most other exhaustion-related genes resulted in distinct grouping of Mitotic, ECs, EM and γδ cells from the remaining subsets. *PDCD1* expression being highest in EM and Mitotic cells, indicating greater intrinsic exhaustion within these subsets. The majority of *PDCD1^+^* cells co-expressed *TIGIT* (**fig. 1f**), and EM and Mitotic cells, but not ECs, were enriched within this PDCD1^+^ TIGIT^+^ population (Fisher’s exact test, **fig. 1g**).

**Figure 1:**
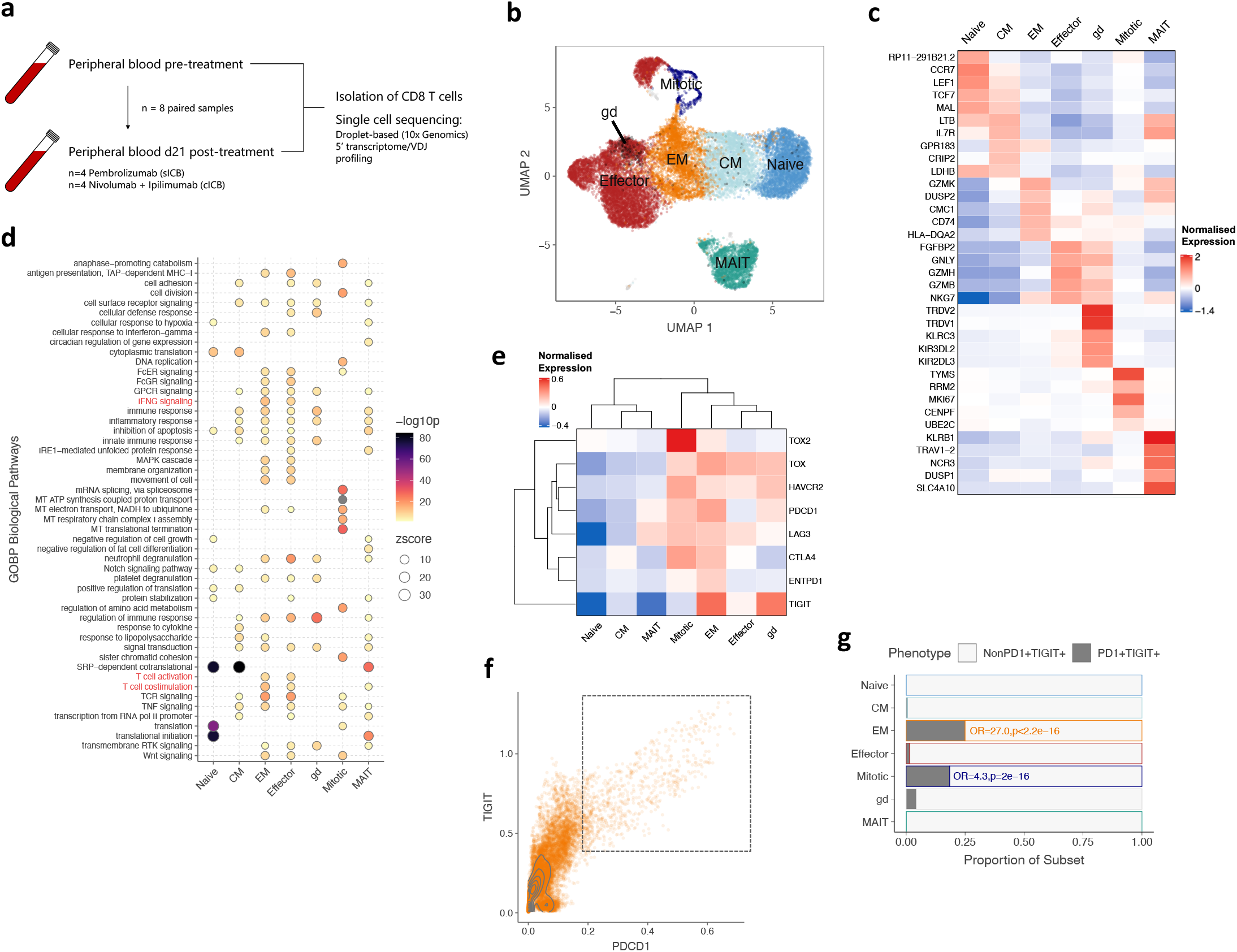
Identification of peripheral CD8^+^ T cell subsets in metastatic melanoma before and during ICB. **(a)** Workflow for sample collection and processing for scRNA-seq. **(b)** UMAP clustering of n=22,445 peripheral CD8^+^ T cells from n=8 individuals sampled at pre-treatment and following 21 days of ICB, across seven distinct subsets as represented by each colour. **(c)** Expression heatmap showing distinct gene expression profiles for the identified subsets. Genes were selected as the five most differentially-expressed genes by p-value present at both pre-treatment and at 21d post-ICB for each timepoint. **(d)** Dot plot heatmap showing Gene Ontology Biological Process (GOBP) terms significantly enriched across each subset (adjusted p-value < 0.05); selected pathways highlighted in red. **(e)** Expression heatmap of selected T cell activation and exhaustion markers. **(f)** Dot plot of imputed *PDCD1* and *TIGIT* expression across CD8^+^ T cells, with the indicated PD1^+^ TIGIT^+^ peripheral exhausted (Ex) population. **(g)** Subset proportions of non-exhausted and exhausted cells, showing an enrichment of EM and mitotic cells by Fisher’s exact test (odds ratio > 1).

### Changes in CD8^+^ T cell subset composition during early immune checkpoint blockade

Focussing on the key subsets (Naïve, CM, EM, EC and Mitotic), we first assessed ICB effects on CD8^+^ T cell compartment composition. We found ICB significantly reduced CM proportion (**fig. 2a**), whilst other subsets remained relatively unchanged. To explore the generalisability of these findings across a larger cohort we developed composite expression signatures for each single-cell subset (**Methods**), characterising them in bulk RNA-sequencing data generated from similarly derived MM patient CD8^+^ T cells*(18)* (n=212 paired C1/C2 samples from 106 individuals). For most single-cell samples (15/16), bulk RNAseq was generated, permitting validation of score-inferred proportions, with significant correlation between bulk expression scores and the single-cell proportions across all subsets (**fig. S2a**). Across the cohort, naïve, CM and mitotic scores anti-correlated with age, whilst EM and EC scores were correlated, in keeping with known ageing effects *(24)* (**fig. S2b**). Notably, whilst ICB reduced naïve and CM expression scores and increased EM and mitotic scores (**fig. 2b**), no significant change in the EC score was observed, indicating the EC subset remains stable. The increase in mitotic score post-ICB remained sustained at C4 (d63) of treatment (**fig. S2c**). Finally, the decrease in naïve and CM score was strongly correlated within individuals, and inversely correlated with mitotic score increase (**fig. S2d**). Changes in EC score correlated with EM, and to a lesser extent, mitotic score, indicating shuttling from these subsets into the mitotic pool, potentially allowing for EC compartment replenishment post-ICB.

**Figure 2.**
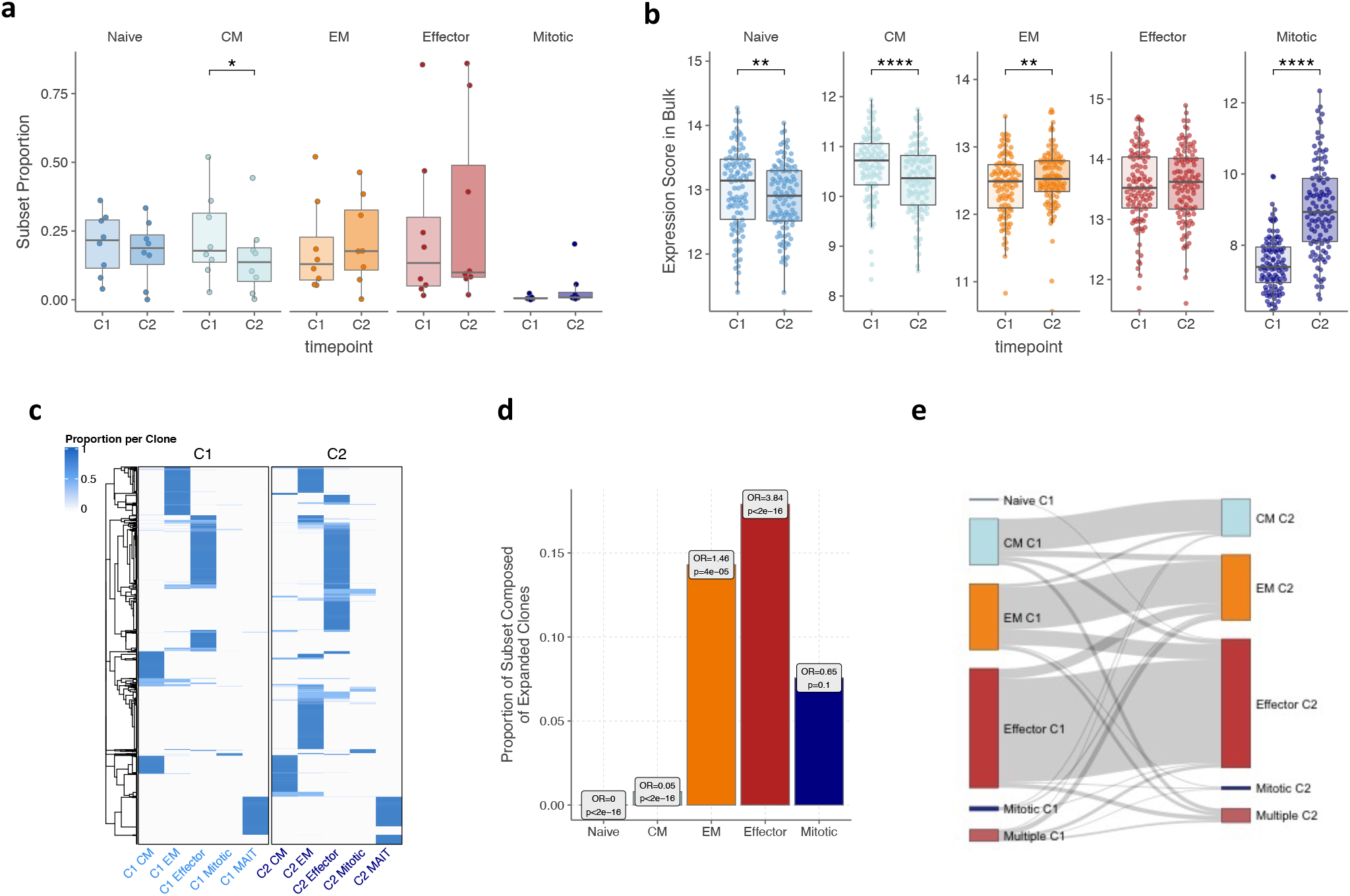
Dynamic immune composition changes upon ICB. **(a-b)** Changes in the proportion of each cell subset between pre-treatment (C1) and 21d post-ICB (C2) as determined by scRNA-seq (n=8, two-sided Wilcoxon-signed rank test) **(a)** or inferred from bulk RNAseq data (n=110, two-sided Wilcoxon-signed rank test) **(b)**. For the bulk RNAseq inference, subset scores were calculated for each bulk RNAseq sample based on a list of markers that distinguished each subset at both pre-treatment and 21d post-ICB in the scRNA-seq data (Methods, Supplementary Fig. 2a-b). **(c)** TCR sharing heatmap between C1 and C2 CD8+ T cell subsets, showing T cell clones consisting of more than 1 detected clone and their relative presence and subset proportions at each timepoint. **(e)** Proportion within each subset at C2 of clones that have significantly expanded from C1 (Fisher’s exact test). **(f)** Sankey plot tracking the cell subset phenotype of C1 clones at C2. For each clone, the subset in which most of its members were present in at a given timepoint was designated as the predominant subset for that clone; ‘multiple’ indicates clones equally distributed across multiple subsets. The width of the connecting flowline represents the number of clones with the respective C1 and C2 subset phenotypes.

We then examined how TCR clones composed of more than one cell are distributed across subsets at C1 and C2 (**fig. 2c**). Most of these clones were of an EC phenotype, being detected in this subset at both timepoints, and whilst there was emergence of novel EM and EC clones at C2, notably few clones were shared across subsets at either timepoint *(14, 22)*. To further characterise the Mitotic cells, we tracked their clonal dynamics between C1 and C2. Interestingly, most Mitotic cells represent small clones, with the median clone size being similar to Naïve and CM cells (**fig. S2e**) and not changing with treatment. Expanded clones were not enriched in the Mitotic subset, and were relatively depleted in Naïve and CM subsets post-treatment (**fig. 2d**, **table S4**). In general, whilst there was limited TCR sharing between memory cells and ECs across timepoints, most resampled clones remained in the same subset over time (**fig. 2e**), the exception being the Mitotic subset where only 2.2% of C2 Mitotic clones were present in any subset at C1. Altogether, the observed changes in subset proportion can be summarised by **(i)** a relative increase in EM cells through both emergence and expansion of pre-existing clones (**fig. 2b, 2c, 1h**), a small proportion differentiating from C1 ECs; **(ii)** stability in the EC compartment size accompanied by changes in clonal dispersion, with expansion of a subset of EC clones (**fig. 2b, 2c**) counterbalanced by reduced median EC clone size (**fig. S2f**); **(iii)** an increase of cells in a mitotic state, the majority of which are small clones not sampled at both timepoints (**fig. 1h, 2b, 2d**), indicating that most Mitotic cells do not establish significant clonal populations; and **(iv)** a relative loss of Naïve and CM cells (**fig. 2b**) following ICB.

### Effector CD8^+^ T cells show increased sensitivity to immune checkpoint blockade

We further characterised responses to ICB between C1 and C2 at a sub-population level, focusing on Naïve, CM, EM and ECs. We controlled for increased power to detect differentially expressed (DE) genes (comparing pre- to post-treatment) in larger subsets by subsampling varying numbers of cells across each subset (analysis bootstrapped 100 times), and used multiple DE analysis pipelines (Methods). Across the four subsets, ECs had more DE genes with treatment irrespective of subsample size (**fig. 3a**). When we used a linear model with an interaction term to control for inter-individual variation we made similar observations (**fig. S3a**). To quantify these patterns of DE, we used a cut-off of n=1100 cells per subsample (**Methods**), and calculated the frequency of genes being detected as significant (adjusted p-value below 0.05) across one hundred bootstraps. Genes such as *IL10RA* and *GZMA* were consistently down- and up-regulated by treatment in all bootstraps in ECs, but not in other subsets (**fig. S3b, table S6**). Comparing the DE genes in ECs versus other subsets, we observed that many were exclusively modulated in this subset (**fig. S3c**), suggesting ICB has a distinct effect on these cells. Subset-wise GOBP analysis of the induced and suppressed genes demonstrated that the most modulated pathways were either conserved pathways represented across all subsets, or specific to ECs (**fig. 3b**). Specifically, pathways restricted to ECs encompassed pro-inflammatory and immune pathways including NFκB, MAPK and TNF signalling (**fig. S3d**), further supporting the profound immune-stimulatory effect of ICB on this subset.

**Figure 3.**
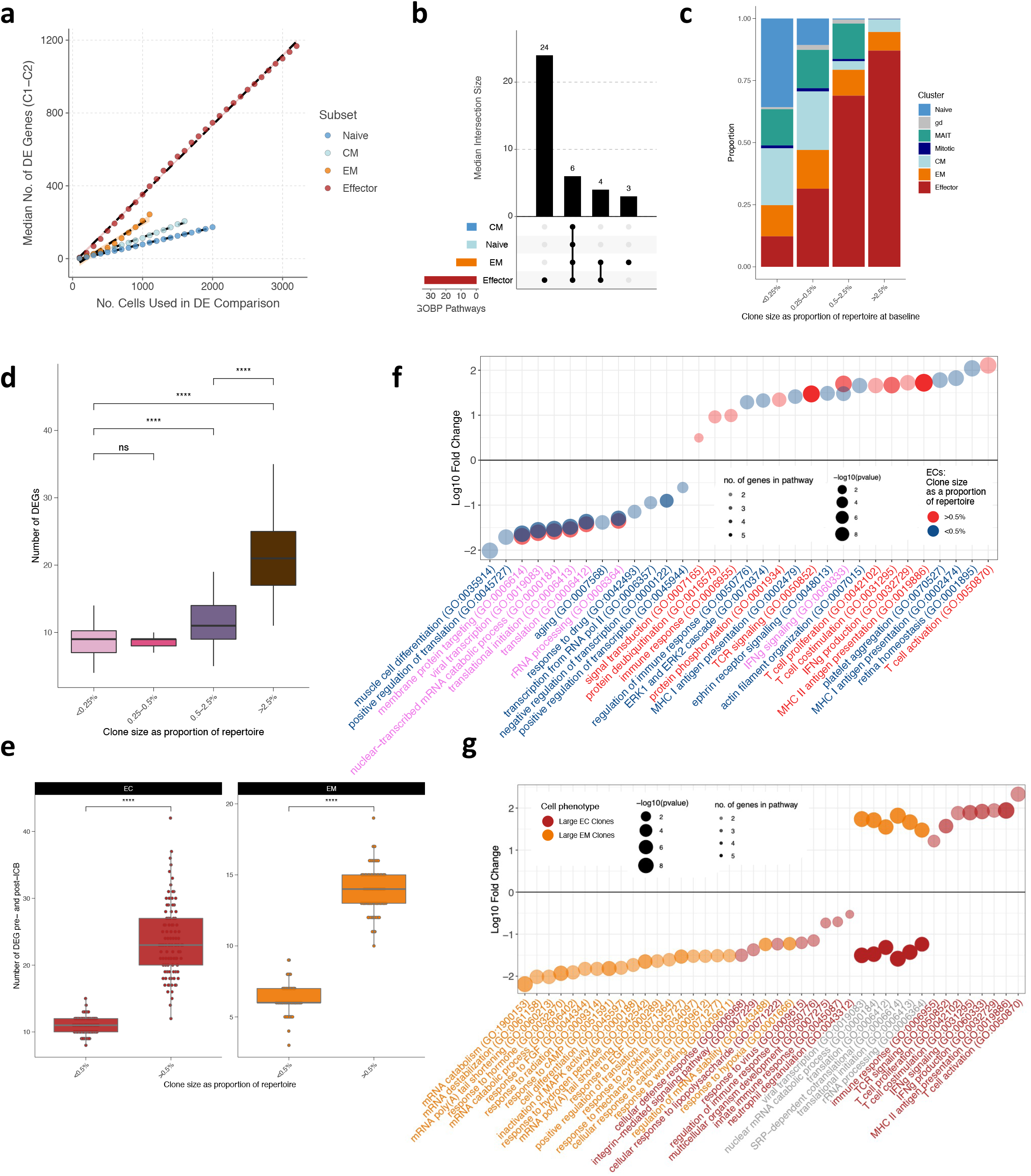
Subset-specific and clone-size dependent transcriptome changes upon ICB. **(a)** Median number of differentially expressed (DE) genes across 21 days of ICB in conventional T cell subsets. Each subset was subsampled to the indicated number of cells at C1 and C2 and tested for DE genes across 100 bootstraps. At each n-value for number of cells, there was a significant difference across the subsets in the number of DE genes (p<0.0001, unpaired Wilcoxon test). **(b)** Upset overlap plot of GOBP pathways upregulated in each subset following treatment (subsample of n=1100 cells), based on the DE genes per bootstrap in (a). The median number of pathways in each intersecting set was calculated over the 100 bootstraps. **(c)** Clonal size bins at C1 filled by proportion of each phenotypic cluster. **(d)** Number of DE genes pre- and post-treatment per clonal size bin. Each clonal size bin was subsampled to a constant size and the analysis bootstrapped 100x. **(e)** Number of differentially expressed genes pre- and post-treatment for large vs small ECs (red) and EMs (orange). Each cluster was subsampled to a constant size and the analysis bootstrapped 100x. **(f)** GOBP analysis demonstrating pathways up- and down-regulated in large vs small ECs pre- and post-treatment. Large ECs are coloured in bright red, small ECs in blue. Pathways where the text is in purple represent those that are shared between both groups. **(g)** GOBP analysis demonstrating pathways up- and down-regulated in large ECs (red) vs large EMs (orange) pre- and post-treatment. Pathways where the text is in grey represent those that are shared between both groups. For all analysis comparing numbers of DE genes, p-values were determined by two-sided Wilcoxon rank-sum test.

### CD8^+^ T cell clones differentially modulate gene expression post-ICB as a function of their size

Given our previous findings linking the number of large clones with clinical outcome, we sought to understand the relationship between the size of a clone and its response to ICB. We grouped clones into four categories according to their size as a proportion of the total repertoire (**fig. 3c**). As expected, most Naïve cells were of the smallest sizes, whereas the larger clone categories were composed of ECs and, to a lesser extent, EMs (**fig. 3c**). We next assessed the relationship between clonal size and ICB effect by performing category-wise DE analysis across treatment (including only clones present at both timepoints, agnostic to subset type), using sampling to control for variation in cell number across sub-population and timepoint, bootstrapping analysis 100 times. This produced a response pattern of DE genes according to clone size, with small and medium-sized clones having comparatively fewer DE genes than large and very large clones (**fig. 3d**).

Given this analysis is potentially confounded by differing subset proportions in each size category, we repeated this process restricted to EC and EM subsets, with clonal size simply dichotomized with a cut-off of 0.5% of the repertoire to denote large from small. The previously observed pattern was preserved within each subset, with larger clones again showing greater numbers of DE genes following ICB compared to smaller clones (Wilcoxon test, p<0.0001, **fig. 3e**). These findings were replicated upon sequential exclusion of each individual in the cohort, suggesting that they are not simply a feature of inter-individual variation (**fig. S3e**).

To understand these effects at a functional level, we analysed GOBP pathways corresponding to the DE gene sets within the EC and EM subsets (**fig. 3f, g; table S7**). We again noted that large EC clones had a strikingly divergent gene expression profile following ICB compared to both small EC clones (**fig. 3f**) and large EM clones (**fig. 3g**), with pathways including T cell activation, proliferation and co-stimulation, TCR signalling, and IFNγ production uniquely upregulated in large ECs.

Taken together, we find clonal size influences both the magnitude of response to ICB as well as the types of genes regulated, and this phenomenon is observed even within the same phenotypic subset. Larger clones display marked enhancement of immune function following ICB treatment, with large ECs uniquely upregulating a number of immune pathways critical to the ICB response.

### Cytotoxic clones demonstrate propensity to persist post-ICB treatment

To further understand the temporal dynamics of clones we tracked the presence or absence of each clone across C1 and C2, restricting the analysis to clones composed of two cells or more. Perhaps unsurprisingly, large clones were more likely to be resampled, with clones only sampled at one timepoint predominantly grouped within smaller size categories (**fig. 4a**). Genes correlated with IFNγ are informative as to cytotoxicity. We therefore adopted previous methods (13) to develop a cytotoxicity score for each cell based on key IFNγ correlated genes. We find that whilst cytotoxicity was globally increased at C2, clones sampled at both timepoints or only at C2 show increased cytotoxicity over those present only at C1 (**fig. 4b**). This finding was robust to separating clones on their size (**fig. S4a**). To explore these observations further we examined baseline (C1) profile of clones, grouping those significantly shrinking (as per a Fisher’s exact test, see **Methods**) or absent at C2 as ‘involuting’ clones. We contrasted these with clones that were found to expand or showing no significant change in size (stable). Interestingly, small clones that subsequently expanded or remained stable following ICB were significantly more cytotoxic at baseline than those that involuted (**fig. 4c**). This finding was similarly observed if we determined change to clone size by simple fold change (**Methods**, **fig. S4b**). This difference in baseline cytotoxicity across involuting and non-involuting clones was not seen within larger clones, which were overall significantly more cytotoxic than smaller clones (**fig. 4c**). As such, these findings suggest that following ICB there is an overall increase in cytotoxicity which is driven by a combination of the persistence of pre-treatment cytotoxic T cell clones, involution of clones with low cytotoxicity and the emergence of more cytotoxic clones.

**Figure 4.**
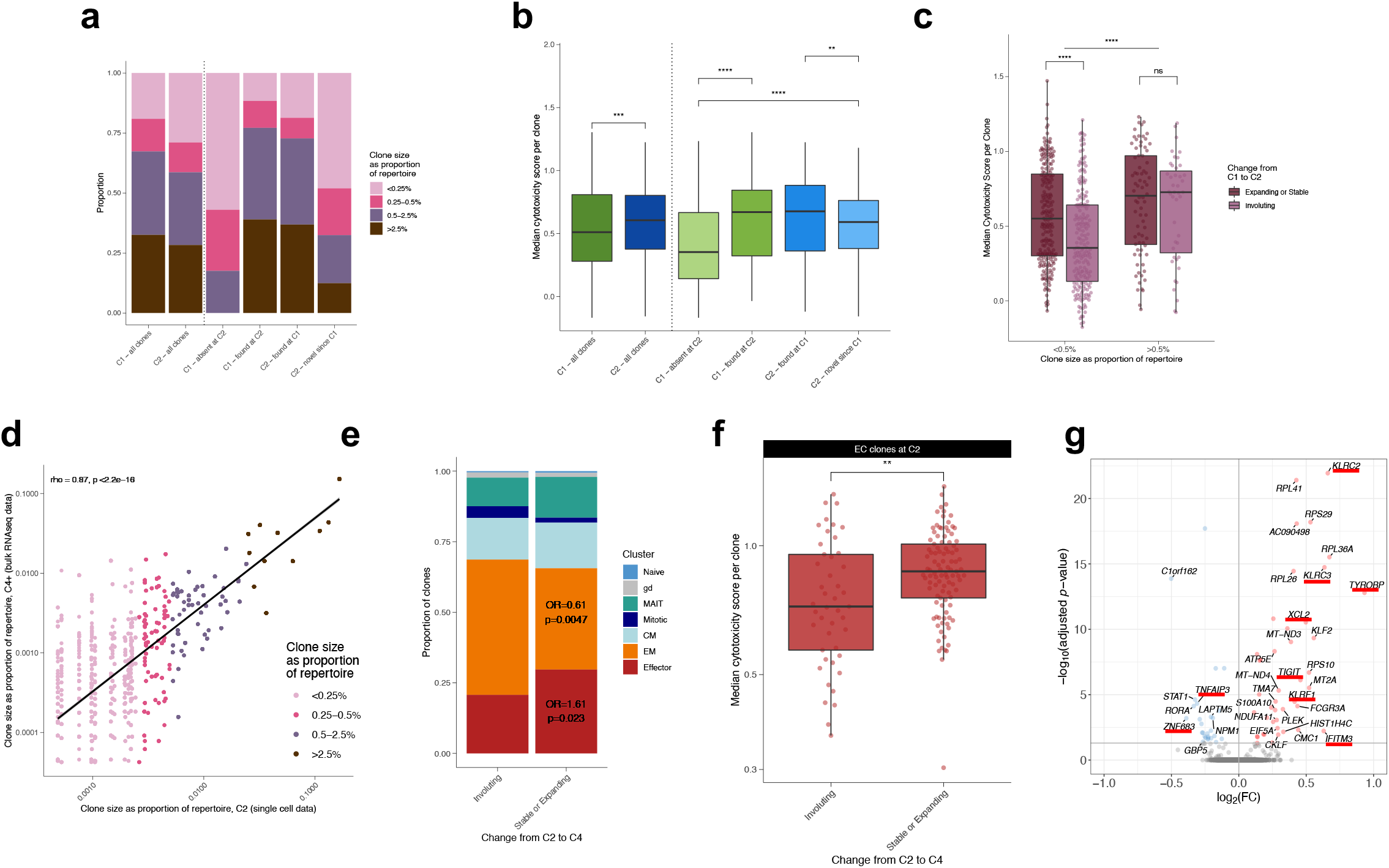
Cytotoxic clones demonstrate propensity to persist post-ICB treatment. **(a-b)** Proportion of clonal size groupings (a) and median cytotoxicity per clone (b) for clones found at both C1 and C2 versus at just at one timepoint, compared to all clones at C1 and C2. **(c)** Median cytotoxicity per clone for those present at C1 that subsequently expand/remain stable (dark purple) or involute (light purple) by C2. Clones are separated based on size using a cut-off of 0.5% of the repertoire to denote large from small. The horizontal line demonstrates the significant difference between all large compared to all small clones. **(d)** Clone size at C2 compared to size at C4+; each point denotes a clone and is coloured based on size grouping at C2 (Spearman’s correlation). **(e)** Proportion of clones that are stable/expanding or involuting from C2 to C4+coloured by their phenotypic cluster; odds ratios for the proportion of ECs and EMs in the expanding/stable compared to the involuting group (Fisher’s exact test). **(f)** Median cytotoxicity score per clone across EC clones found at C2 that subsequently expand/remain stable or involute by C4+. **(g)** Volcano plot of genes differentially expressed by ECs at C2 that subsequently expand/remain stable by C4+ compared to those that involute; genes of interest are denoted with an underscore. The direction is in favour of clones that expand/remain stable – i.e., those in red with a positive FC are upregulated in clones that expand/remain stable. For all analyses, only clones with >1 cells were included, with unpaired Wilcoxon tests used for all comparisons unless otherwise stated.

To extend analysis across later timepoints, we examined bulk RNA-seq data from CD8^+^ T cells isolated 63-106 days post-treatment (C4+), available for six out of the eight individuals for which scRNA-seq data was available (**Methods**). From the corresponding TCRs identified in these samples, we found 992 overlapping unique-TCRβs associated with 4,245 cells at C2 within the scRNA-seq data. Notably, clone size at C2 and C4+ were highly correlated (Spearman’s rho=0.84, p<2.2×10^-16^, **fig. 4d**), indicating relative preservation of clonal structure over this longer period. Comparison of C2 clones (minimally 2 cells in size) that expanded or remained stable versus those that involuted between C2 and C4+ highlighted an enrichment for ECs in the stable/expanding group (OR 1.6, p=0.023) whilst there was under-representation of EMs (OR 0.61, p=0.0047) (**fig. 4e**). We subsequently constrained further analysis to the EC subset, analysing C2 cytotoxicity of stable/expanding clones versus those involuting by C4+. Again we found a significantly higher cytotoxicity in those stable or expanding (**fig. 4f**), and this was robust to alternatively defining changes to clone size based on fold change, rather than by a Fisher’s test (**fig. S4c**). Notably, this trend persisted when clones were grouped based on their size (**fig. S4d**), although was not statistically significant for the large clones, likely reflecting underpowering. Finally, by comparing expression of EC clones stable/expanding between C2 to C4+ to those that involuted we identify genes linked to clonal stability. Intriguingly, we find killer lectin receptor genes including *KLRF1* (Killer Cell Lectin Like Receptor F1), *KLRC2/3*, and the marker *TIGIT*, were amongst those significantly higher in stable/expanding ECs at C2, whilst downregulated genes included *STAT1* **(fig. 4g, table S8**), suggesting these may form markers of clonal persistence.

Collectively, these results indicate that pre-existing cytotoxicity belies increased clonal persistence, with increased C1-C2 persistence of clones that are more cytotoxic at baseline, whilst emerging clones, although small, also possess higher levels of cytotoxicity. Similarly, within the EC compartment, C2 cytotoxic clones have increased likelihood to be found at d63+ of treatment.

### Baseline and early on-treatment cytotoxicity is prognostic for clinical outcome during ICB

Given that baseline cytotoxicity delineates cells likely to persist or expand during ICB, potentially reflecting response to antigen, we explored the relationship between both pre and on-treatment CD8^+^ T cell cytotoxicity and clinical response to ICB. We generated cytotoxicity scores for each bulk sequencing sample in our cohort (n=131 at C1 and n=109 at C2, **Methods**) which we separately validated with flow cytometry across a subset of samples, finding strong correlation of the cytotoxicity score with expression of cytotoxic markers such as perforin-1 (**fig. S5a-b**). Whilst cytotoxicity at C1 was strongly-correlated with C2 cytotoxicity across the cohort, indicative of the importance of the baseline response (**fig. S5c**), ICB tended to increase cytotoxicity, with a larger effect from cICB **(fig. S5d**).

Patients had significantly higher CD8^+^ cytotoxicity pre-treatment compared to healthy controls (**fig. 5a**), reflecting the systemic immune effects of MM. Of note, in patients who proceeded to respond to treatment, both C1 and C2 cytotoxicity was higher than in those progressing by six months. Using Kaplan-Meier estimates (log-rank test), we found a cytotoxicity score above the cohort median at either baseline or C2 construed increased progression free survival (PFS) (**fig. 5b**, **S5e**), supporting the finding that baseline immune profile is important in determining clinical outcome, and demonstrating that agnostic sampling of the peripheral CD8^+^ subset provides prognostic information. In contrast, we found no association between the mitotic score and PFS at either C1 or C2 (p=0.78 and 0.69 respectively, log-rank test, data not shown), suggesting the degree of baseline CD8+ T cell division, or that induced by ICB, does not relate to clinical benefit.

**Figure 5.**
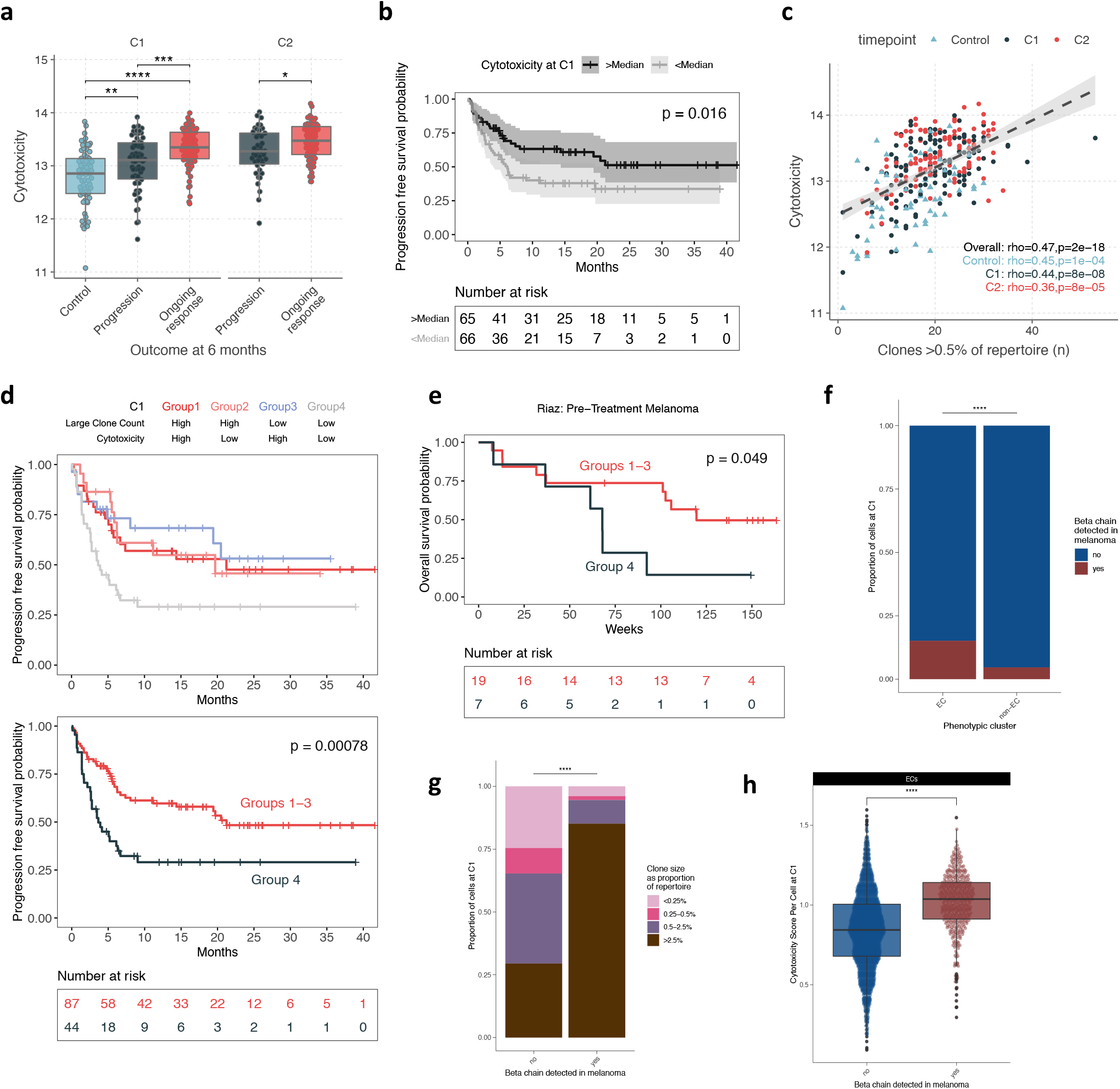
CD8^+^ T cell cytotoxicity is associated with clinical response to ICB. **(a)** Cytotoxicity calculated in CD8^+^ bulk RNA-seq samples is significantly higher in responding patients than progressor and control individuals at pre-treatment and d21 (n=69 controls, 112 patients). **(b)** Kaplan-Meier curves showing increased progression free survival in patients with baseline cytotoxicity above median (n=131 patients) (two-sided log-rank test). **(c)** Correlation between large clone count and cytotoxicity score in the bulk cohort, across control individuals and patients at C1 and C2 (n=69 controls, 137 patients, Spearman’s rank test). **(d)** Progression free survival in patients separated by a combination of C1 large clone count and cytotoxicity (above/below median) (n= 131, two-sided log-rank test; Group1 vs Group4, p=0.019; Group2 vs Group4, p=0.023; Group3 vs Group4, p=0.010). **(e)** Analysis of overall survival in melanoma patients from Riaz et al. 2017*(25)* based on intratumoral RNAseq and TCRB-seq data. Patients were separated by a combination of pre-treatment large clone count (TCRB-seq) and cytotoxicity (RNAseq) (above/below median, as in (d)) (n=26, two-sided log-rank test). **(f)** Proportion of ECs compared to non-ECs at C1 that carry a tumour-infiltrating TCRB chain, as identified by TCRseq of resected melanomas by Pruessman et al. 2020*(26)* (Fisher’s exact test). **(g)** ECs at C1 that are either tumour-infiltrating or not, filled by their clonal size; Fisher’s exact test for the proportion of large clones (>0.5%) compared to small clones found in the tumour-infiltrating and not-tumour-infiltrating compartments. **(h)** Cytotoxicity scores of ECs at C1 categorised by whether they carry a tumour-infiltrating TCRB chain or not (unpaired Wilcoxon test).

Having previously shown that the number of large clones at C2 is associated with clinical response to ICB*(18)*, we explored whether CD8^+^ cytotoxicity could be integrated with large clone count to stratify patients according to outcome *prior* to treatment. First, we recapitulated in this expanded cohort the finding that C2 but not C1 large clone count is strongly associated with PFS, with increased significance in the cohort as it expands and matures (**fig. S5f**). As expected, CD8^+^ cytotoxicity score correlated with large clone count across healthy individuals, C1 and C2 samples (**fig. 5c**), whilst responding individuals had significantly higher cytotoxicity irrespective of large clone count when assessed with a linear model (**fig. S5g**). We next explored whether cytotoxicity and large clone count could additively predict six-month patient outcome using a linear discriminant analysis (LDA) model. This yielded an area under the receiver operating characteristic curve (AUC) of 0.69 (**fig. S5h**), but did not seem to be co-operatively affected by cytotoxicity or large clone count (**fig. S5i**). Instead, given the close association between the two measures, we wondered whether they had a compensatory interaction. We grouped individuals on a combination of above/below median large clone count and cytotoxicity and examined their PFS on this basis. We found that individuals with below both median large clone count *and* median cytotoxicity (Group 4) at baseline had significantly lower PFS compared to all other groups (**fig. 5d**) which otherwise had comparable survival; this effect also being observed at d21 (**fig. S5j**). As such, we further stratified a subset of patients with low pre-treatment peripheral CD8^+^ T cell activity, with limited clonal expansion and low cytotoxicity, that receives minimal clinical benefit from treatment with ICB.

### Validation using external datasets

In order to establish the validity of our findings and apply them more generally, we sought relevant published datasets. To explore the prognostic value of cytotoxicity score in an alternative context we looked to see whether our observations extended to the tumour environment. Utilising the Riaz dataset of paired RNA-seq and TCRB-seq data from melanomas prior to and during treatment with anti-PD-1*(25)*, we devised an intratumoral CD8^+^ T cell cytotoxicity score by selecting genes used in our peripheral cytotoxicity analyses that showed strongest correlation with *CD3D, CD3E, CD8A, CD8B, IFNG* and *PRF1* expression (**fig. S6a**), and subsequently identified the number of TCRB clones per sample with proportion above 0.5%. Despite the low number of individuals in this cohort, we found significantly lower overall survival for patients with below median large clone count and cytotoxicity (**fig. 5e**) at baseline.

Given this concordance between the peripheral and intratumoral T cell state, we used an alternative dataset of TCRB chains sequenced from melanoma biopsies *(26)*. From the 199 samples, 85,528 unique TCRB were identified, of which 392 were found in our samples. We assessed the characteristics of cells carrying one of these known tumour-infiltrating beta chains at both timepoints and found these contained a significantly increased proportion of ECs compared to other cells subsets (**fig. 5f, S6b**). Within these ‘TIL-TCR’ ECs, there was additionally a significantly higher proportion of large clones (**fig. 5g, S6d**) with higher cytotoxicity score at both timepoints (**fig. 5h, S6c**). Extending these observations to our bulk cohort, we also observed greater clonal expansion of TIL-TCR matching T cell clones in patients compared to healthy controls (**fig. S6e**). Notably, when we compared this to the profile of cells matching public viral-reactive TCRs *(27)* (n=149 cells), clonally expanded ECs were under-represented, and there was no significant difference in cytotoxicity (**fig. S6f-h**). Together, this supports our findings of the importance of large, peripheral EC clones which appear to be over-represented within the tumour-infiltrating compartment.

## DISCUSSION

Identifying the cells most responsive to ICB is vital in understanding the mechanisms of ICB response and resistance. In this study we have addressed the peripheral CD8^+^ T cell response to ICB at a single-cell level, characterising responses by sub-populations as well as by clones. We identify seven phenotypic subsets that, as well as sharing a conserved response to ICB, display subset-specific changes, the greatest and most divergent being within the effector cells (ECs). The T effector memory (EM) subset has been described as a crucial mediator in light of its expansion intratumorally and peripherally*(15, 28, 29)*, and we observe increases in the peripheral EM score, inferring commensurate expansion of this subset. We show though that, whilst ICB does not change EC proportion, the EC compartment exhibits the greatest magnitude of clonal and gene expression changes and preferentially persists post-ICB. As such, we posit that in peripheral blood, the effector T cell compartment is most sensitive to ICB treatment. In keeping with this, recent work shows that clones common to blood and tumour are enriched for effector markers and are not peripherally exhausted in patients who respond to ICB*(14)*.

That ICB preferentially affects large clones reveals mechanistic underpinnings to our previous observations linking large peripheral CD8^+^ T cell clone count with clinical outcome*(18)*. Smaller clones downregulate expression of immunological pathways and increase expression of ribosomal genes and translational activity post-ICB; whereas, consistent with the observation that translational suppression occurs in terminal effector T cells, larger clones have an opposite response*(30)*. It is unclear to what degree clonal size is a causal factor in the differential response to ICB or correlates with other factors. However, that large EC clones show relative baseline upregulation of TCR signalling, likely reflective of successful TCR ligation and subsequent expansion, suggests the increased sensitivity to ICB reflects intrinsic activity of these clones.

The observation that ICB is associated with survival and expansion of clones that display markers of cytotoxicity *prior* to treatment was supported by the inclusion of on treatment bulk-sequencing data collated at later cycles of treatment. This highlighted that cytotoxic clones persist across multiple cycles of ICB, consistent with previous descriptions of a relationship between cytotoxicity and response to PD-1 blockade in the context of NSCLC*(31)*. Although it is difficult to definitively prove that ICB specifically elicits clonal survival, we find that ICB does lead to increased clonal cytotoxicity. Given persistent clones and expanding clones demonstrate higher cytotoxicity pre-treatment or at C2, these data support this to be an action of ICB peripherally. This potentially also implies limits to patient response to ICB centre on deficient tumour immunogenicity or antigen presentation, with consequential limited effector T cell responses*(32)*. Thus therapies targeted at overcoming further immune checkpoints may have relatively diminished clinical returns compared to those improving antigen presentation and priming.

Consistent with the literature, we find that ICB provides a marked stimulus to mitotic cell division in CD8^+^ T cells. The relevance of this response to outcome is less clear however. Notably, the Mitotic subset showed increased expression of exhaustion markers, and most mitotic clones did not appear to become established at C2 or at later timepoints, with only a small proportion of surveyed mitotic CD8^+^ T cells progressing to EC and EM cells. Whilst this implies dysfunction, whether these are classically exhausted or are in state with features of exhaustion is unclear and will form the basis for future research. Conversely, we observe that ECs remain persistently expanded for long durations, and our work suggests Killer-Lectin Receptors may specifically denote long-lived ECs.

From a clinical outcome perspective, our work further demonstrates that analysis of peripheral CD8^+^ T cells both prior to and during treatment provides prognostic information. The association of large clone count with response to ICB likely reflects the increased probability an individual will carry tumour reactive clones as this count rises. Nonetheless, a situation where a few specific anti-cancer clones are present could be envisaged to be beneficial, but not register in total large clone count. These cells would anticipated to be cytotoxic though and here we show that a high cytotoxicity score may compensate for low large clone count. Comparison of data from healthy individuals indicates that, not only do healthy individuals have fewer large circulating clones, but they have diminished cytotoxicity. Finally we find no association between the magnitude of the mitotic response and clinical outcome, highlighting a disconnect between mitotic responses and those that are cytotoxic and therapeutically important.

A key question has been how these peripheral clones may relate to tumour - are they bystanders or is there evidence of enrichment with those that react to melanoma antigens? By integration of our results with TCR sequencing of melanoma resident T cells from Pruessman and colleagues*(26)* we demonstrate that public clones found in melanomas are predominantly large, and that such reactive clones show higher than average cytotoxicity. Conversely, although clones carrying TCR reactive to viral antigens are plentiful, they are not larger or more cytotoxic. These findings are consistent with recent descriptions of tumour-infiltrating clones which tend to have an effector-like phenotype in the peripheral blood in mouse models*(17)* and patients with MM*(16)*.

A potential limitation of this study is the inability to definitively causally link the changes observed to ICB exposure - and exclude for example effects of cancer progression or other temporal factors. This is a draw-back of this type of observational work in humans – it is difficult (and ethically challenging) to obtain samples from untreated individuals with cancer over time. Nonetheless, the changes we observe in patient profiles coupled to the frequently pronounced transformations in clinical performance strongly argue that these are due to the effect of ICB. This is underlined by bulk RNAseq data where we observe a greater magnitude of change in cytotoxicity score induced by cICB versus sICB (**fig. S5c**), consistent with an effect of ICB on cytotoxicity. Further, baseline cytotoxicity is higher in melanoma patients than healthy volunteers, and is higher in those that respond to treatment compared to those that do not (**fig. 5a**), suggesting an important interaction between baseline immune state, the effect of ICB on cytotoxicity and response to treatment.

In summary, our work combines in-depth analysis of peripheral CD8^+^ T cells responses to ICB at the single-cell level, with observations corroborated and extended in a large dataset. The use of V(D)J sequencing allows us to track clones over time, determining that most mitotic cells fail to expand into differentiated large effector clones. Instead, clone size plays a key role in determining the magnitude and nature of response to ICB, and we find the effector subset is most sensitive to treatment. Finally, we reveal that cytotoxic clones have successfully expanded to form persistent subsets – and that much of the effect of ICB appears to rely upon this pre-existent response, implicating failure of immune recognition as being the major limiter of response to ICB. Going forward, efforts to understand the durability and maintenance of tumour reactive clones over time will be important. Similarly, understanding the degree to which other, potentially modifiable, non-tumour related factors impact on the ICB response to cancer will be vital in further improving therapeutic outcomes.

## MATERIALS AND METHODS

### Study design

The aim of this study was to understand peripheral CD8^+^ T cell responses to ICB at a single-cell level, in particular, identifying key phenotypic or clonal features of response to therapy. Eight patients who were prescribed ICB for MM at a tertiary Cancer Centre in the UK were prospectively and sequentially recruited. Four participants received single agent Pembrolizumab (sICB) whilst four received combination Ipilimumab/Nivolumab (cICB). Characteristics of the participants are outlined in **table S1.** All participants provided written informed consent to donate samples to the Oxford Radcliffe Biobank (Oxford Centre for Histopathology Research ethical approval nos. 16/A019, 18/A064, 19/A114). 30 – 50 ml blood was collected into EDTA tubes (BD vacutainer system) taken immediately pre-treatment at d0 and d21 following one cycle of treatment (**fig. 1a**). Following single cell isolation, transcriptome and V(D)J sequencing was undertaken with downstream analysis performed in R, as detailed below.

### Sample collection

Peripheral blood mononuclear cells were immediately obtained from whole blood by density centrifugation (Ficoll Paque). CD8^+^ cell isolation was carried out by positive selection (Miltenyi) according to the manufacturer’s instructions, with all steps performed either at 4 °C or on ice. Unless otherwise stated, patients receiving cICB (ipilimumab plus nivolumab, n=4) or sICB (pembrolizumab, n=4) were pooled for the purposes of analysis, based on previous work showing qualitatively similar genetic changes induced in peripheral CD8^+^ T cells by both cICB and sICB*(18)*.

### Single-cell sample preparation and sequencing

Magnetically separated CD8^+^ cells were immediately oil-partitioned into single-cell droplets, followed by cell lysis and a reverse transcription reaction using the 10x Genomics Chromium system. 6000 CD8^+^ T cells were loaded onto each partitioning cassette. Single-cell 5’ RNA transcriptome and V(D)J Libraries were constructed from the cDNA, according to the manufacturer’s instructions (https://www.10xgenomics.com/). Due to the sequential and prospective nature of sample collection, droplet generation and RT steps were performed on individual samples, but library generation and sequencing was performed in two batches of pooled samples (table S1). Sequencing was performed on an Illumina HiSeq4000; 75bp PE reads for the 5’ RNA libraries, 150 bp PE reads for the V(D)J libraries to a depth of approximately 50000 reads per cell.

### Data processing, quality control and clustering

Single-cell reads were aligned, counted and filtered for initial quality control (QC) as previously described*(12)*. In short, FASTQ files were generated from Illumina BCL outputs, used to produce gene expression and barcoding libraries, and then combined using the Cellranger *mkfastq*, *count* and *aggr* functions respectively. For V(D)J data, Cellranger *vdj* was applied. For QC, the R package *scater(33)* was used to identify single-cell outliers as low-quality libraries and *scran* utilised to detect doublets which were removed from analysis. Following alignment and data pre-processing, CD8^+^ T cells were filtered on detectable *CD3D* and either *CD8A* or *CD8B* UMIs whilst lacking *CD14* expression. The R package *Seurat(34)* was used for further QC, data normalisation and identification of variable features; low quality cells were excluded if they had less than 500 total UMIs, less than 300 detected features and mitochondrial gene percentage above 20%. Genes expressed in <5 cells were also removed. The percentage of mitochondrial genes and cell cycle scores were regressed out upon scaling of the normalised data. Principal component analysis (PCA) was performed on the scaled data and the top 16 principal components selected for dimensionality reduction and UMAP clustering as the most variable and biologically informative components. To determine an ideal resolution value, cell clusters generated using values between 0.1 and 2.5 were compared using the R package *clustree(35)* with values between 1.5 and 2.0 showing high cluster stability. Batch was not observed to have impact on Principal Component distribution and there was no significant difference in baseline cytotoxicity score between batches.*(13)*. In short, FASTQ files were generated from Illumina BCL outputs, used to produce gene expression and barcoding libraries, and then combined using the Cellranger *mkfastq*, *count* and *aggr* functions respectively. For V(D)J data, Cellranger *vdj* was applied. For QC, the R package *scater(33)* was used to identify single-cell outliers as low-quality libraries and *scran* utilised to detect doublets which were removed from analysis. Following alignment and data pre-processing, CD8^+^ T cells were filtered on detectable *CD3D* and either *CD8A* or *CD8B* UMIs whilst lacking *CD14* expression. The R package *Seurat(34)* was used for further QC, data normalisation and identification of variable features; low quality cells were excluded if they had less than 500 total UMIs, less than 300 detected features and mitochondrial gene percentage above 20%. Genes expressed in <5 cells were also removed. The percentage of mitochondrial genes and cell cycle scores were regressed out upon scaling of the normalised data. Principal component analysis (PCA) was performed on the scaled data and the top 16 principal components selected for dimensionality reduction and UMAP clustering as the most variable and biologically informative components. To determine an ideal resolution value, cell clusters generated using values between 0.1 and 2.5 were compared using the R package *clustree(35)* with values between 1.5 and 2.0 showing high cluster stability. There was no observed effect of batch on PC distribution or baseline cytotoxicity score.

### Identification of cell subsets

To assign each cell cluster generated in Seurat to a functional T cell subset, we curated transcripts per million (TPM) normalised counts for CD8^+^ T cells from previous T cell cancer scRNA-seq datasets*(12, 13, 36–38)*. Using the R package *SingleR(39)*, each individual cell was matched to one of the CD8^+^ T cell subsets present in the reference datasets based on global gene expression similarity, from which subset identity for each cluster was determined. Briefly, we noted cluster 12 (ECs) was grouped above the Mitotic clusters despite being labelled as EC through *SingleR* and having high expression of FGFBP2 and other EC-characteristic genes; we detected a high level of expression of ribosomal genes in this subset relative to other ECs, explaining its projection on the UMAP despite being an EC cluster.*(13, 14, 39–41)*. Using the R package *SingleR(39)*, each individual cell was matched to one of the CD8^+^ T cell subsets present in the reference datasets based on global gene expression similarity, from which subset identity for each cluster was determined.

### Clonal definitions, size and emergence vs involution

T cell chains were filtered based on called productivity, and chain identity (TRA or TRB). Cells were selected as those that either contained a single TCRβ chain, a combination of one TCRα with one TCRβ chain or a group of three chains – two TCRα and TCRβ. Cells not meeting this criteria were excluded. A clone ID was defined by a concatenation of amino acid CDR3 chains present, with cells sharing identical clone IDs classed as members of the same clonotype. Clonal proportions were calculated by dividing the number of cells in each clone per sample by the total of number of cells (meeting the above criteria) per sample. Out of 22446 cells which had transcriptome sequencing that had passed QC, 17909 had matching V(D)J sequencing with required chain combinations as specified above, with these cells used for all clonal analyses.

For analyses examining subset phenotype of clones in terms of size and evolution (e.g., **fig. 2e, S2f**), cells were grouped by clone ID, individual and timepoint, and proportion of its member cells within each subset was calculated; the highest proportion subset was used to describe the clone’s major subset. For global overview of clonal size at C1/C2 within each subset, clonal size was defined depending on the timepoint the clone was sampled at. Analysis of changes in subset identity per clone (**fig. 2d**) utilised clones present at both timepoints only.

For analyses looking at clonal changes across timepoints, only clones present at both timepoints were included (e.g., **fig. 3d-g**). In this context, the size of the clone was defined based on its C1 (pre-treatment) size. For clones that are present at only one timepoint (e.g., **fig. 4a**), the clone size was defined based on its size when present. Similarly for analyses only at one timepoint (e.g., **fig. 4c-g, 5f-h**) clonal size was defined based on size at that timepoint.

For the analysis of expanding/stable versus involuting clones between C1 and C2, only clones with >1 member cell were included and only cells present at C1 were analysed (**fig. 4c**). For clones that were found at both timepoints, clones were designated to be involuting if the proportion at C2 was significantly (a nominal p value <0.05, Fisher’s exact test) smaller than the proportion at C1 and vice versa for expanding. All clones that were found at C1 (with >1 member cell) but not found at C2 were also classed as involuting. Expanding cells were grouped with those in which there was no significant difference between timepoints (p>0.05, Fisher’s exact test) as the ‘stable or expanding’ subset. We also performed an additional analysis where clones were classified as involuting based on a range of fold changes to size (**fig. S4b**). Here, ‘involuting’ clones were defined as those where the proportion at C2 was either <0.6x or <0.4x the proportion at C1 and ‘stable or expanding’ clones were all others.

For the analysis of expanding/stable versus involuting clones between C2 and C4+, again only clones with >1 member cell were included and only clones present at C2 were analysed. Clones present at C4+ were identified based on the presence of the TCRB amino acid sequence within bulk RNAseq data of magnetically sorted CD8^+^ T cells for six of the same individuals. A clone was designated as being present at C2 and C4+ if that same TCRB was found at both timepoints (C2 in scRNA-seq, C4+ in bulk RNAseq). The clonal size (as a proportion) at C4+ was calculated by diving the number of reads for that TCRB by the total number of TCR reads in the sample. The designation of a clone into involution or expanding/stable followed the same methodology as described above, including a sensitivity analysis using FC cut-offs.

### Downstream expression analysis

#### Differential gene expression analysis with subsampling

Differential gene expression was performed in *Seurat* using the FindMarkers function for all genes present, unless otherwise indicated. Alternatively, to validate the FindMarkers results, a mixed linear model for all genes with variance >0 in the subsampled cells was used, controlling for interindividual variation (**fig. S3a, d**). Where specified, to control for differences in statistical power in detecting DE genes across groups with varying cell counts, the number of cells used for analysis was standardised to the number present in the smallest group and bootstrapped 100 times to account for sampling bias. For comparison of DE genes according to subset or clone size, 1100 or 245 cells were subsampled respectively as the minimum number of cells across each category at both timepoints. This same methodology was repeated filtering for only ECs or EMs with cells grouped into large and small clones based on a 0.5% of repertoire cut-off and subsampling performed to the smallest group size.

#### Pathway analysis

Pathway enrichment analysis was performed using the R package *XGR(40)* with the GOBP database. Significantly induced or suppressed genes were assessed separately against a background of all detected genes using a hypergeometric test, ontology algorithm specified as ‘elimination’ and an elimination p value of 0.01. For comparisons of pathways modulated following ICB across subsets, pathways were assessed for DE gene from each subsample at n=1100 cells, and the frequency of enriched pathways was used for comparison; with this approach employed to maximise the number of DE genes/statistical power taken forward for pathway analysis in most subsets given the paucity of DE genes in Naive cells. For comparisons of pathways modulated following ICB in large versus small ECs/EMs, we included the minimum number of significant DE genes across all comparators, keeping the number of genes used in the analysis constant between groups.

#### Leave-one-out cross validation

In order to explore the effect of inter-individual variation, we repeated the differential gene expression with subsampling analysis eight times, with all cells from one individual excluded each time. This demonstrated consistency in results suggesting that the observed effect is not driven solely by cells from one donor.

#### Generation of cytotoxicity score

Cell scores for cytotoxicity were calculated using AddModuleScore in Seurat, based on the top 50 variable feature genes that significantly correlated with *IFNG* expression *(41)* (see supplementary data file 2). For PDCD1 and TIGIT expression analysis, the MAGIC R package was used to recover expression data*(42)*. In most instances – and as indicated in the figures and figure legends – cytotoxicity score is reported on a clonal basis. Here, cells were grouped for each individual and timepoint based on clone ID and the cytotoxicity score calculated as a median of the cytotoxicity scores for all cells within that clone.

### Bulk RNA-sequencing

Bulk RNA-sequencing was performed on CD8^+^ T cells from 69 healthy individuals and 140 patients receiving single or combination ICB for MM, with exclusion of adjuvant treated patients. Out of 137 C1 and 113 C2 samples, 110 individuals had paired C1/C2 data with a further 49 having C4 samples. Clinical follow-up was available for 132 C1 and 109 C2 samples. Gene expression and TCR analysis using the MiXCR package was performed as previously described; for full methods, please *see(18)*.Data can be downloaded from the European Bioinformatics Institute and the Centre for Genomic Regulation under accession no. <u>EGAS00001004081</u>.

To generate subset scores from each bulk sample, hallmark genes for each subset were identified in the single-cell data using the FindAllMarkers function in Seurat across all cells from C1 and C2 samples separately. To minimise timepoint-specific genes, the top 20 genes at each timepoint were selected and the overlapping genes retained as robust subset-demarcating genes (**table S2**). Scores were then generated for each gene-set using *DESeg2*-normalised expression data *(43)* to then calculate geometric mean expression for each group of subset genes. A similar approach was used for cytotoxic score but instead based on the top 50 genes used for single-cell cytotoxic scoring. All metadata is listed in **table S11**.

For clonal persistence at C4+, we used longitudinal bulk samples from six individuals for which single-cell data was obtained; four taken at day 63, one at day 84 and one at day 106 of treatment. Clones were defined within these samples using MiXCR *(44)* according to the beta chain, and then compared with earlier scRNA-seq data with matching beta chain usage.

### Flow cytometry

Patient PBMCs frozen in 90% FCS + 10% DMSO were thawed at 37°C and washed once in HBSS prior to staining of 1×10^6^ cells with LIVE/DEAD Fixable Near-IR (Invitrogen). Cells were washed and subsequently stained with antibodies against CD3, CD8, CD16, CD56, CD45RA, CD27, CX3CR1 and HLA-DR in 5% FBS + HBSS. After washing, cells were fixed for 10 minutes with 2% paraformaldehyde (Sigma), washed and then permeabilised with 1x Permeabilisation Buffer (eBioscience) for 10 minutes. The cells were then stained with Perforin-1, CCL4, Granzyme-B, IFN-g, TNF-a, IL-32 and Granulysin, washed with perm buffer and stained with anti-rat IgG2a secondary, before a final wash with perm buffer and acquisition on a BD LSRFortessa X-20. Data was analysed in FlowJo (FlowJo LLC). Antibodies details are shown in **table S9** and gating is outlined in **table S10**. All staining steps were performed for 30 minutes and at 4°C unless otherwise stated.

### Analysis of external datasets

Melanoma tumour TCRB-seq and FKPM-normalised RNA-seq from Riaz et al. 2017 *(25)* were downloaded from https://github.com/riazn/bms038_analysis or from GEO:GSE91061 respectively. To generate an intratumoral cytotoxicity score, the 50 genes used for single-cell cytotoxic scoring were correlated against the expression of *CD3D*, *CD3E*, *CD8A*, *CD8B*, *IFNG* and *PRF1* and k-means clustered to identify a module of genes most associated with CD8^+^ T cell cytotoxicity. Geometric mean expression of the eight genes was then calculated per sample and used as cytotoxic score. Large clone count per TCRB sample was calculated as for the bulk CD8 sequencing data (i.e., the number of clones occupying greater than 0.5% of the repertoire). Survival analysis was conducted for the 52 samples with overlapping RNA-seq and TCRB data (26 pre-treatment, 26 on-treatment) (**table S12**). Analysis of public melanoma and viral T cell clones was done using TCRB-seq data of 199 melanomas from Pruessman et al. 2020 *(26)*, or viral-reactive TCR sequences downloaded from VDJdb *(27, 45)* [04/2021] with confidence scores above 0. Cells present in the scRNA-seq data were annotated as being found in melanoma or viral-reactive based on their ownership of a TCRB that was found in either the Pruessman or VDJdb dataset.

### General statistical analysis

Statistical comparisons were performed as indicated in each figure and conducted in R. Paired tests were used for paired samples. Cells and statistics from donor 6 were excluded from all analyses in figure 2c-d, S2e and figure 4 due to low numbers of cells at C1. For analysis of clonal persistence (**fig. 4d-g**), n=6 as only six out of the eight individuals profiled had samples at C4+. All analyses and statistical tests were performed in R using baseR or *rstatix*, and *lme4/lmerTest* for linear effect models. LDA analysis was performed using *MASS* and *Pi (46)*. Gene correlation analysis for phenotype scores was performed using *DGCA(47)*. Survival analysis was performed using *survival/survminer. UpSetR(48) networkD3* and *ComplexHeatmap(49)* were used for visualisation. For all data, *p<0.05, **p<0.01, ***p< 0.001, ****p<0.0001. Adjusted p-values have been reported using Bonferroni correction for multiple comparisons.

## LIST OF SUPPLEMENTARY MATERIALS

Supplementary tables are found in excel workbook as separate tabs

S1: Single Cell Characteristics

S2: Subset Scoring

S3: Cytotoxic Scoring

S4: TCR information

S5: Movement of clonal subsets

S6: Subset DEGs

S7: Subset + clone DEGs

S8: Stable vs involuting clone markers

S9: Flow Cytometry Antibodies

S10: Flow Cytometry gating

S11: Bulk RNAseq Metadata

S12: Riaz et al. Analysis

## ACKNOWLEDGEMENTS

We are very grateful to all patients who generously contributed samples and participated in the study. We thank all the staff of the Day Treatment Unit, Oxford Cancer Centre, and The Brodey Centre at the Horton General Hospital. We also thank M. Payne, N. Coupe and R. Matin for assistance in collecting patient samples. We thank M. H. Al-Mossawi for discussion and advice.

## AUTHOR CONTRIBUTIONS

RAW & BPF conceived the study; RAW, OT, IN & BPF performed the analysis; figures were generated by OT & RAW; manuscript was drafted by RAW, OT & BPF; RAW & RC performed single-cell experiments; OT and PKS performed flow cytometry experiments; RAW, EAM, RC, CAT, PKS, AV & OT collected and prepared samples; HR provided expert statistical advice; MRM contributed to discussions throughout the work; BPF oversaw the project. All authors read, edited and approved the final manuscript.

## DATA AND MATERIALS AVAILABILITY

Raw expression matrices and associated cell metadata for all cells that passed upstream QC will be deposited on EGA. All scripts used in data analysis and the generation of figures will be made available on the Fairfax group bitbucket account.

## SUPPLEMENTARY FIGURES

**Supplementary Figure 1:**
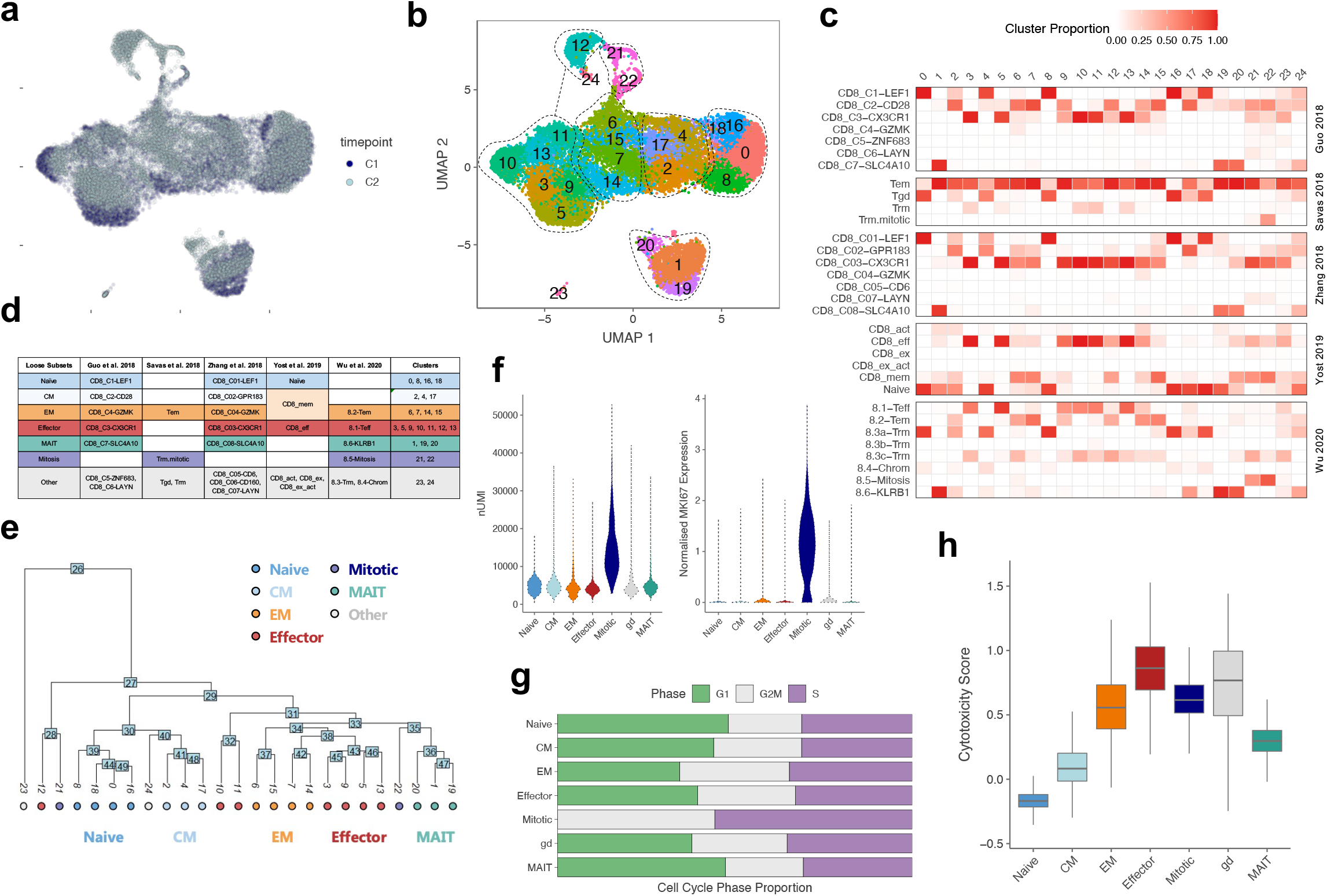
Workflow for CD8^+^ T cell subset assignments. **(a)** UMAP distribution of cells from pre-treatment (C1) and day 21 on-treatment (C2) samples. **(b)** UMAP visualisation of 25 Seurat clusters generated at a FindClusters resolution of 1.5. **(c)** Heatmap of assigned subset identities for each Seurat cluster using external cancer patient single-cell RNAseq datasets as a reference*(12, 13, 36–38)*. For each cell, SingleR was used to cross-match its expression profile to that of one of the subsets in each reference set. The proportion of cells in each cluster that were assigned a particular identity from each reference set is represented by the intensity of shading (between 0 and 1). **(d)** Summary table of subset identities for each reference set and assigned identities for each cluster based on the best match from the SingleR analysis. **(e)** Phylogenetic tree for each Seurat cluster coloured by assigned subset showing similarity between clusters within the same subset. **(f)** Violin plot showing the nUMI (left) and normalised MKI67 expression (right) per cell across each of the seven CD8^+^ T cell subsets. **(g)** Bar plot of the proportion of cells in each stage of the cell cycle, assigned using the CellCycleScoring function in Seurat, across each subset. **(h)** Cytotoxicity score (based on expression of the 50 most *IFNG*-correlating genes, Supplementary Table S3) across each subset.

**Supplementary Figure 2:**
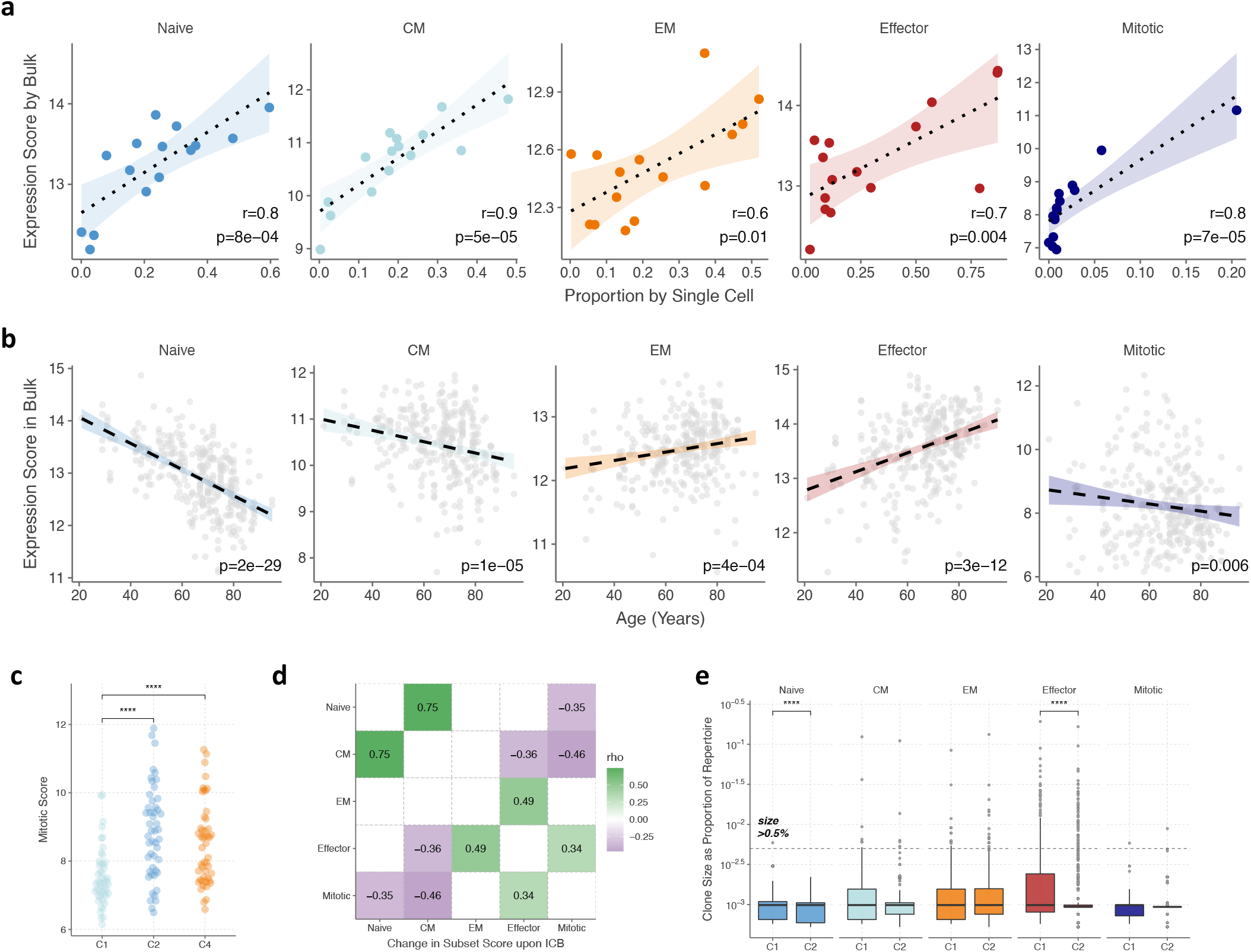
Inference of subset proportions and phenotype. **(a)** Correlation between subset proportions for each single-cell sample and scores calculated for each subset using bulk RNAseq data of CD8^+^ T cells. For 15/16 of the samples used for single-cell RNAseq, bulk RNAseq on the purified CD8^+^ T cells was also performed. A list of the top 20 discriminating markers for each subset was generated using FindAllMarkers in the single-cell data for all pre-treatment cells, and separately for all cells at d21 post-ICB. The overlapping genes were retained as markers of each cell subset irrespective of treatment, and the geometric mean expression for each set of genes was calculated using normalised bulk RNAseq data, thus generating subset scores for each bulk RNAseq sample. Pearson’s correlation was used to determine the accuracy of this scoring approach for each subset, showing good concordance for all subsets. **(b)** As a further validation of the bulk RNAseq scores, subset scores across n=399 samples were correlated against participant age using a mixed linear model, controlling for cycle as a fixed variable; lmer(Score ~ Age+(1|Cycle)). **(c)** Changes in mitotic scores at C1 (d0), C2 (d21) or C4 (d63) of treatment (n=49 individuals, two-sided Wilcoxon-signed rank test). **(d)** Heatmap of Spearman’s rho values for correlations between bulk subset scores; filled boxes represent adjusted p-value <0.05 upon Bonferroni’s correction. **(e)** Change in clonal size with major phenotype in the given subset at C1 and C2 (unpaired Wilcoxon test). For each clone, the subset in which most of its members were present was designated as the predominant phenotype for that clone; clones equally distributed across multiple subsets were excluded. Clones above the dotted line represent large clones occupying size greater than 0.5% of the repertoire.

**Supplementary Figure 3.**
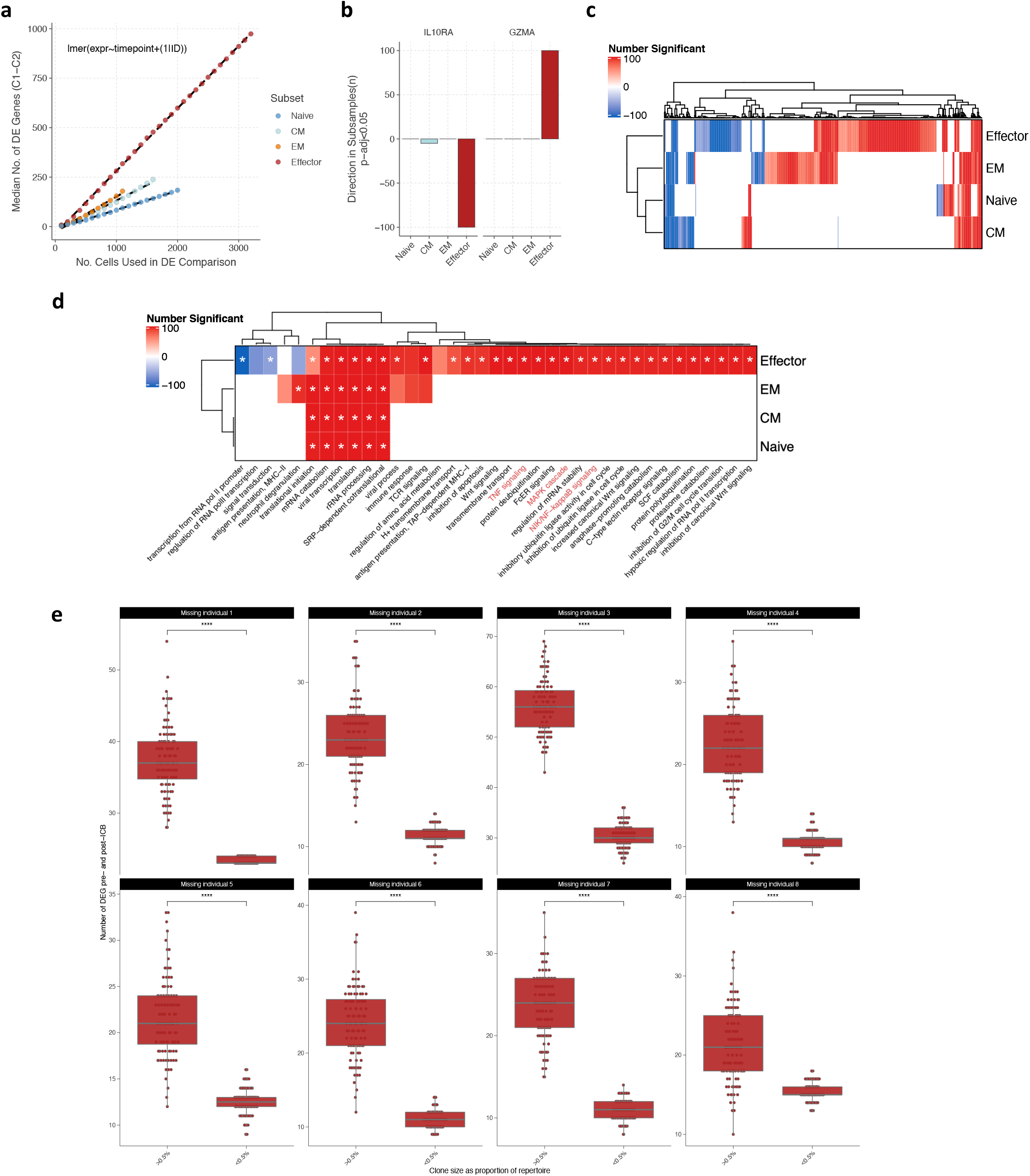
Differential expression analysis across subsets and clone sizes. **(a)** Median number of differentially expressed (DE) genes across 21 days of ICB in conventional T cell subsets, as per Fig 3a but using the annotated mixed linear model instead of the *FindMarkers* Seurat function to identify significantly modulated genes (Bonferroni’s adjusted p-value <0.05). At each n-value for number of cells, there was a significant difference across the subsets in the number of DE genes (p<0.0001, unpaired Wilcoxon test). **(b)** Differential modulation of *IL10RA* and *GZMA* across each subset using *FindMarkers* bootstraps. The number of times each of the specified genes was detected as significantly up- or downregulated across 100 bootstraps was plotted per subset (subsample of n=1100 cells, direction represents increased or decreased expression following ICB). **(c)** Heatmap of DE genes across each subset, detected in greater than 50/100 *FindMarkers* bootstraps. Intensity of shading indicates how many bootstraps the gene was detected in, and colour represents whether the gene was significantly up- or downregulated. **(d)** Heatmap of induced or suppressed GOBP pathways modulated in each subset, in more than 50/100 *FindMarkers* bootstraps; selected pathways highlighted in red. Pathways marked with an asterisk* represent those also found using an equivalent approach based on DE genes detected by the mixed linear model. **(e**) Number of differentially expressed genes pre- and post-treatment for large vs small ECs (red) sequentially omitting one individual each time. The EC cluster was subsampled to a constant size and the analysis bootstrapped 100x (unpaired Wilcoxon test).

**Supplementary Figure 4.**
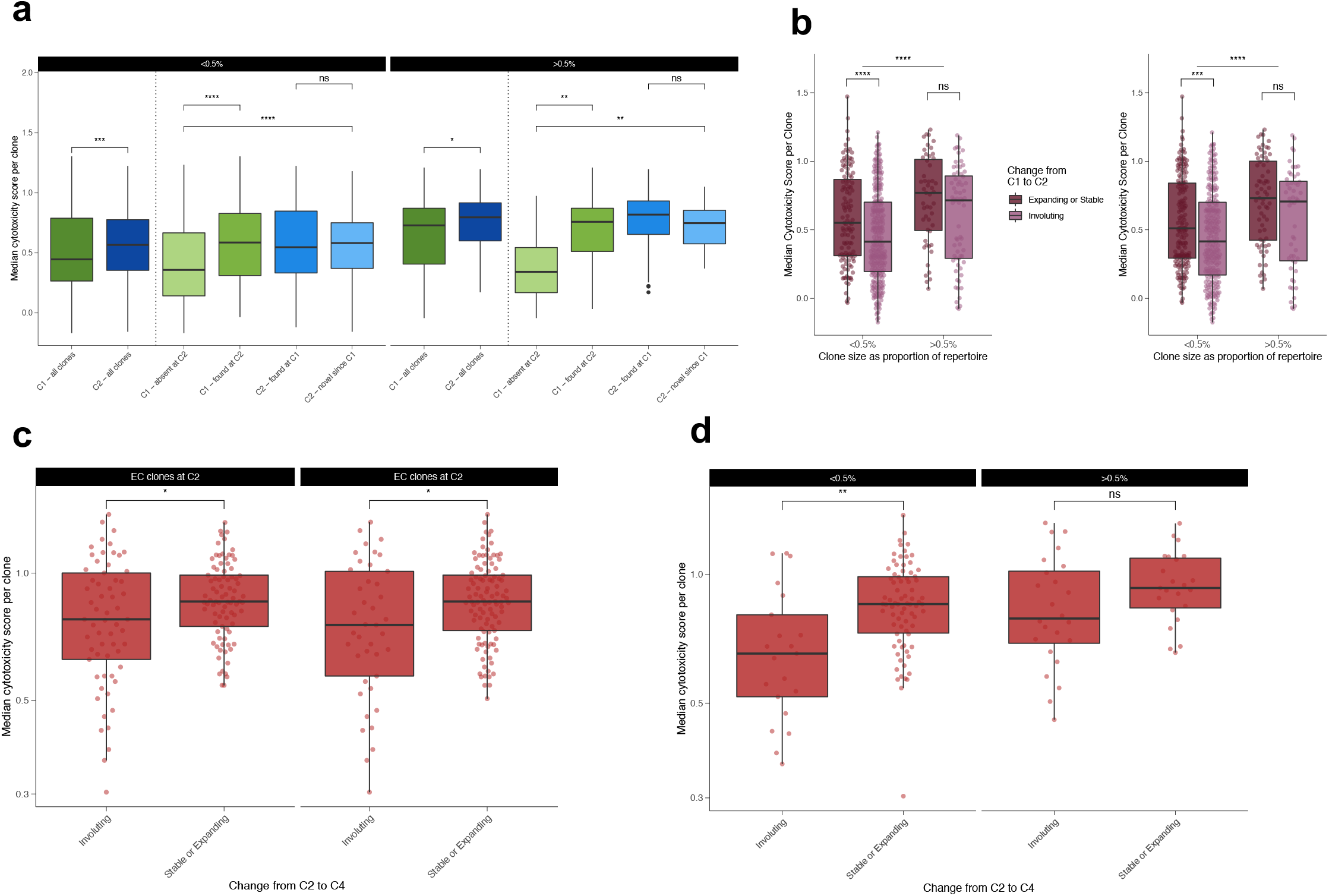
Cytotoxic clones demonstrate propensity to persist post-ICB treatment. **(a)** Median cytotoxicity per clone for those found at both C1/C2 versus just at one timepoint, relative to all clones at C1 and C2. Clones were separated based on clonal size using a cut-off of 0.5% of the repertoire to denote large from small. **(b)** Median cytotoxicity per clone for those present at C1 that subsequently expand/remain stable (dark purple) or involute (light purple) by C2. Clones are separated based on size using a cut-off of 0.5% of the repertoire to denote large from small. Here, involuting clones are defined as those which are either 40% (left panel) or 60% (right panel) smaller at C2 compared to C1, whilst expanding/stable clones are those which, at C2, are within 40% or 60% of their baseline size. **(c)** Median cytotoxicity score per clone across EC clones found at C2 that subsequently expand/remain stable or involute by C4+. Here, involuting clones are defined as those which are either 40% (left panel or 60% (right panel) smaller at C4+ compared to C2, whilst expanding/stable clones are those which, at C4+, are within 40% or 60% of their C2 size. **(d)** Median cytotoxicity per clone for EC clones found at C2 that subsequently expand/remain stable or involute by C4+, separated based on clonal size using a cut-off of 0.5% of the repertoire to denote large from small. For all data, Wilcoxon rank-sum tests used for all comparisons unless otherwise stated.

**Supplementary Figure 5.**
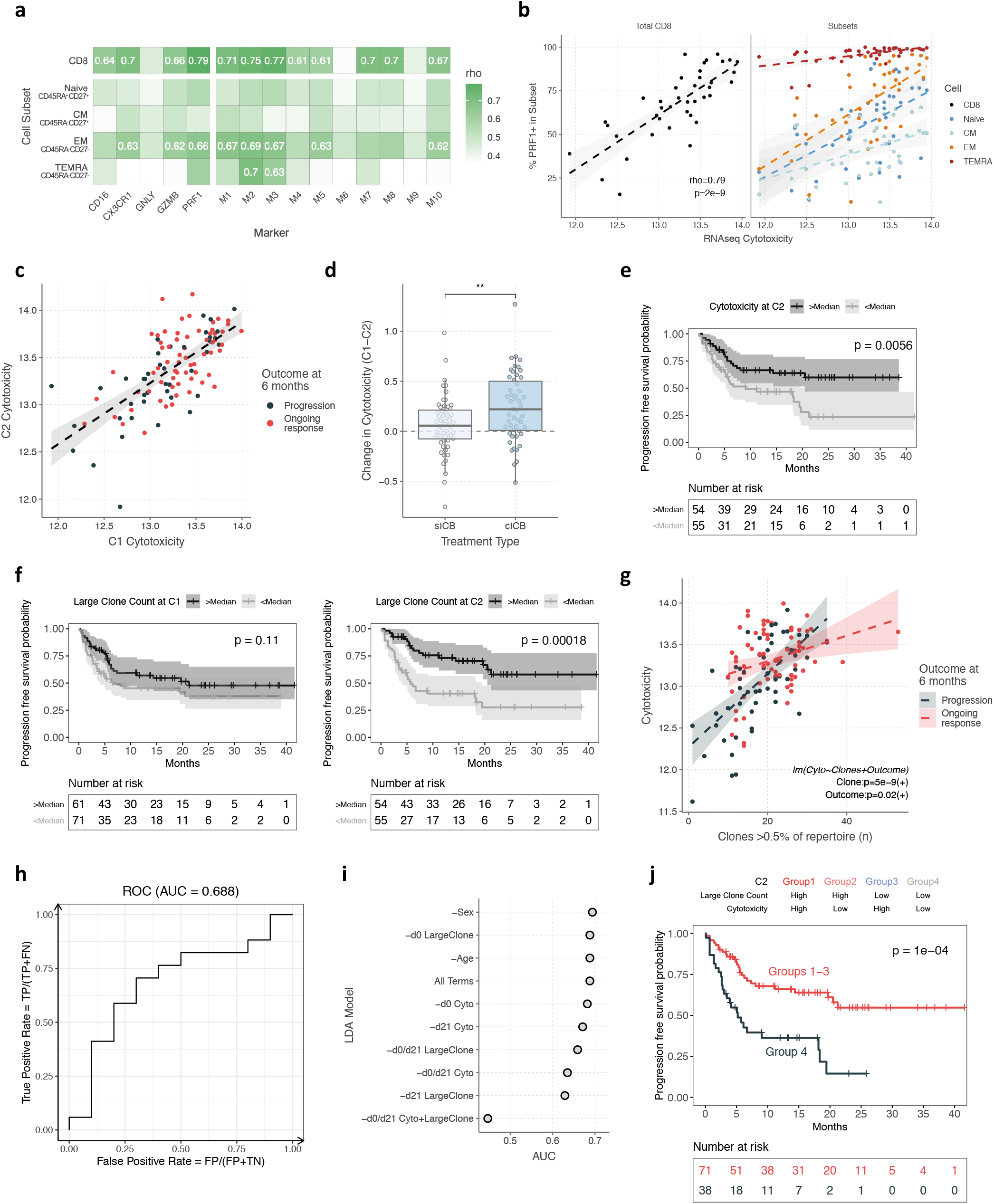
CD8^+^ cytotoxicity associations with flow cytometry, TCR analysis and clinical outcomes. **(a)** Heatmap ot Spearman’s correlations between bulk RNA-seq cytotoxicity and cytotoxic protein expression by flow cytometry across all CD8 T cells, or within CD8 cell subsets (n=20 C1 and 20 C2 samples; gating indicated on axis). M1-M10 refers to combinatorial gates of cytotoxic proteins; all gating listed in Supplementary Table 9. The top 10 Spearman’s rho values are displayed in the heatmap. **(b)** Spearman’s correlation between bulk RNA-seq cytotoxicity and % PRF1^+^ cells across all CD8^+^ T cells (left) or within subsets (right). **(c)** Correlation between cytotoxicity at C1 and C2 (n=106 patients, Spearman’s rank test). **(d)** Change in cytotoxicity in bulk cohort between C1-C2 based on ICB treatment type (n=106 patients, two-sided unpaired Wilcoxon-test). **(e-f)** Kaplan-Meier curve of progression-free survival in patients with d21 cytotoxicity **(e)**, or d0 (left) and d21 (right) large clone count **(f)** above median above median (n=132 patients or 109 patients at d0 or d21 respectively, two-sided log-rank test). **(g)** Linear effect model testing for correlation between C1 cytotoxicity, and large clone count and six-month clinical outcome, controlling for both as covariates (n=132 patients, positivity or negativity of estimate value indicated in brackets). **(h)** Receiver operating characteristic plot for a linear discriminant analysis (LDA) model for six-month clinical outcome, incorporating d0 and d21 large clone count and cytotoxicity, age and sex (n=106 patients). Individuals were randomly separated into independent training (n=78) and test (n=28) sets for cross-validation. **(i)** Area under curve (AUC) values for the LDA model in (h) upon omitting various predictor variables. **(j)** Progression free survival in patients separated by a combination of d21 large clone count and cytotoxicity (above/below median) (n=109, two-sided log-rank test).

**Supplementary Figure 6.**
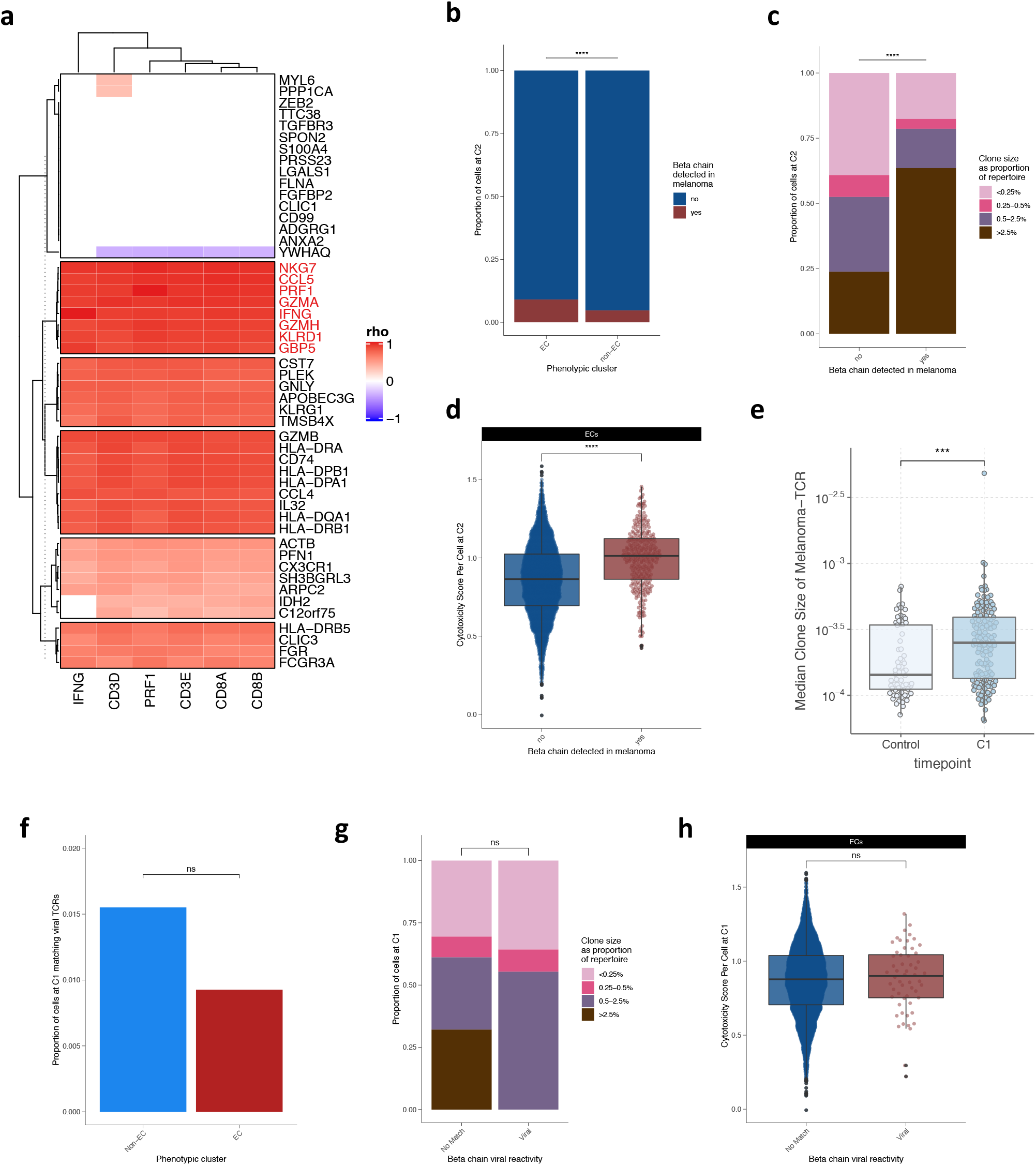
Intratumoral analysis of T cell clonality and cytotoxicity. **(a)** Heatmap of Spearman’s correlations based on tumour RNAseq data from Riaz et al. 2017 between *CD3D, CD3E, CD8A, CD8B, IFNG* and *PRF1* gene expression and 50 genes used in the calculation of peripheral cytotoxicity score, with the selected module of most-correlated genes used in the calculation of intratumoral T cell catotoxicity highlighted in red. **(b)** Proportion of ECs compared to non-ECs at C2 that carry a tumour-infiltrating TCRB chain, as identified by TCRseq of resected melanomas by Pruessman et al. 2020*(26)* (Fisher’s exact test). **(c)** ECs at C2 that are either tumour-infiltrating or not, separated by clonal size; Fisher’s exact test for the proportion of large clones (>0.5%) compared to small clones found in the tumour-infiltrating and not-tumour-infiltrating compartments. **(d)** Cytotoxicity scores of ECs at C2 categorised by whether they carry a tumour-infiltrating TCRB chain or not (unpaired Wilcoxon test). **(e)** TCRB chains were matched with the TCR sequences in the bulk cohort and the median clone size of matching clones per individual was compared between healthy controls and melanoma patients (unpaired Wilcoxon test). **(f-h)** TCR chains from the single-cell VDJ data were matched with viral TCR sequences from VDJdb and the relative subset enrichment **(f**; Fisher’s exact test), clonal size groupings in ECs **(g**; Fisher’s exact test for proportion of clones >0.5% across groups) and cellular cytotoxicity in ECs **(h**; unpaired Wilcoxon test) was similarly compared between viral-matching and non-matching cells.

